# Growing neuronal islands on multi-electrode arrays using an Accurate Positioning-µCP device

**DOI:** 10.1101/026401

**Authors:** Robert Samhaber, Manuel Schottdorf, Ahmed El Hady, Kai Bröking, Andreas Daus, Christiane Thielemann, Walter Stühmer, Fred Wolf

**Author notes:** equally contributing authors (Robert Samhaber, Manuel Schottdorf, Ahmed El Hady).

## Abstract

**Background:** Multi-electrode arrays (MEAs) allow non-invasive multiunit recording in-vitro from cultured neuronal networks. For sufficient neuronal growth and adhesion on such MEAs, substrate preparation is required. Plating of dissociated neurons on a uniformly prepared MEA’s surface results in the formation of spatially extended random networks with substantial inter-sample variability. Such cultures are not optimally suited to study the relationship between defined structure and dynamics in neuronal networks. To overcome these shortcomings, neurons can be cultured with pre-defined topology by spatially structured surface modification. Spatially structuring a MEA surface accurately and reproducibly with the equipment of a typical cell-culture laboratory is challenging.

**New Method:** In this paper, we present a novel approach utilizing microcontact printing (μCP) combined with a custom-made device to accurately position patterns on MEAs with high precision. We call this technique AP-μCP (accurate positioning micro-contact printing).

**Comparison with existing Methods:** Other approaches presented in the literature using μCP for patterning either relied on facilities or techniques not readily available in a standard cell culture laboratory, or they did not specify means of precise pattern positioning.

**Conclusion:** Here we present a relatively simple device for reproducible and precise patterning in a standard cell-culture laboratory setting. The patterned neuronal islands on MEAs provide a basis for high throughput electrophysiology to study the dynamics of single neurons and neuronal networks.

## INTRODUCTION

The culturing and survival of living cells in vitro requires the preparation of suitable conditions in an artificial environment. In particular, in order to study the electrophysiological properties of dissociated neurons, a close contact between the cells and the recording electrodes has to be established. The negative surface charge and hydrophobic nature of unmodified glass surfaces are known to counteract attachment and growth of neurons. One way to modify the surface properties in a favourable way is by coating with growth and adhesion promoting molecules to allow attachment, development and cell survival. Multi-electrode arrays (MEAs) offer a versatile and well-established tool for both, non-invasively studying activity patterns in neuronal networks on a wide range of spatial scales (Gross et al., 1977, 1995, 1997; Stett et al., 2003; Hofmann et al., 2006, 2011; Schottdorf et al. 2012; Potter et al., 2011) and biosensor applications (Keefer et al., 2001; Chiappalone et al., 2003; Selinger et al., 2004; Martinoia et al., 2005; Xiang et al., 2007).

MEAs are devices in which a thin layer of a conducting material in the form of an electrode array is embedded onto the surface of a glass substrate, allowing for non-invasive parallel recording and stimulation of electrical activity at multiple sites from a cell culture. Plating dissociated neurons on a uniformly coated culture substrate results in random network formation that are highly variable in their detailed topology. A way to align the growth of cell processes with the predefined topology of the MEA and to reduce inter-culture variability is to apply patterned substrate modifications that are aligned with the predefined topology of the MEA layout.

Culturing neurons on MEAs for extracellular stimulation and signal recording, together with the ability to precisely and reliably pattern neuronal networks, is thus a crucial step in the development of neuroelectronic hybrids such as biosensors, neuronal prostheses and neuroelectronic circuits.

Patterning neurons in a predefined topology requires that geometric parameters like pattern layout, dimension and alignment to a substrate can be adjusted reproducibly and precisely. Several methods to achieve a predefined topology in cultured neuronal networks have been proposed in the past: topographical-patterning and chemical-patterning. Pioneered in the 1960s, different topographical patterning techniques included etching groves on a substrate and lithographic procedures to directly model three dimensional features on a culture substrate (Niemeyer et al., 2004). Chemical patterning methods include patterned deposition of adhesion promoting proteins (Wheeler et al., 1999; Scholl et al., 2000; Nam et al., 2004). Using the cell repelling properties of Polyethylenglycol (PEG) through a photolithographic process was also shown to effectively direct cell growth (Kumar et al., 1994). All of these methods, however, require specialised equipment not readily available in a standard cell-culture laboratory setting. Additionally, the alignment of the pattern to be cultured with a given substrate layout requires additional high-precision equipment. More recently, seminal work on patterning using carbon nanotubes has been performed by the group of Yael Hanein (Shein-Idelson M et al. 2011) where islands of carbon nanotubes were deposited on the electrodes leading to the preferential growth of clusters of neurons over the electrodes. One-dimensional neuronal cultures, which provide a platform to study the propagation speed of neuronal signals, have been realized on multielectrode array using a combination of adhesive and protein repelling coating (Jacobi S, Moses E 2007). Glial islands on which monolayers of neurons or single neurons are grown have been widely used in electrophysiological studies (Lau M, Bi G 2005, Burgalossi et al. 2012) and have provided a means to pattern neuronal cultures. Micro-contact printing (μCP) through patterned Polydimethylsiloxan (PDMS) micro stamps allows for a relatively simple approach to transfer adhesion promoting molecules such as Laminin or Poly-D-Lysine to substrate surfaces at high spatial resolution (Wheeler et al., 1999; Lauer et al., 2001; James et al., 2004; Chang et al., 2006; Jun et al., 2007). While this method does require a photo-lithography laboratory in order to produce a positive template for the stamps, continued access is not required once the stamp has been produced.

In the current method paper we present a simple positioning device and micro-contact printing technique (accurate positioning micro-contact printing, AP-μCP). To characterize the utility of the AP-μCP procedure, we designed an island-pattern fitting the electrode layout of a MEA. Circular patterns were established by micro-contact printing the adhesion promoter on the MEA. The islands were aligned to the electrodes, allowing for the growth of isolated populations of neurons. We show that the AP-μCP technique yields reproducible and topologically defined neuronal islands arbitrarily aligned with the electrodes of a MEA. We also patterned neurons in a one dimensional geometry allowing electrophysiological measurement of activity propagation through a one dimensional neuronal culture.

Our study thus presents a simple, precise and reliable patterning technique that can serve as an elementary but crucial component for throughput electrophysiology of single neurons and neuronal networks.

## MATERIALS AND METHODS

### 3.1 master production through photo-lithography

According to the designed pattern, a chrome coated soda-lime mask was produced by electron beam lithography (ML&C, Jena). Photoresist layers (AZ 9260; Microchemicals, Ulm, Germany) of 20 μm were spin-coated on glass wafers and subsequently soft-baked at 100°C for 12 min. Structuring of the photoresist-layer was obtained by exposure to UV irradiation for 12 minutes in close contact with the mask carrying the negative pattern. With proper care and handling, no wear on the masters could be observed during the course of this study. In this study, we used three predesigned patterns, two which realized “Islands” patterns of 64 islands of either 90 μm or 60 μm diameter each separated by 200 μm (Fig. 3A). The second pattern was designed as a “Highway” pattern (Fig. 7A) of 100 μm in width and 2 mm in length (Fig. 7A)

### 3.2 pdms preparation

PDMS preparation was performed by mixing the PDMS Silicone Elastomer Base and Curing Agent (Sylgard ^®^ 184; Dow Corning, Wiesbaden Germany) in a proportion of 10:1. The prepared volume was stirred vigorously by hand and subsequently degassed under a vacuum bell jar: The lidless volume containing the mixed PDMS agents was placed in the bell jar and evacuated with a vacuum pump. After 3 minutes, the pump was turned off and the bell jar left in low pressure (100-300 hPA) for 20 minutes to allow all air bubbles to escape. The PDMS exhibits effervescence during evacuation. Slower application of low pressure prevented this.

### 3.3 the mould

The stamps were cast from PDMS into a stainless steel mould, the interior surfaces of which had been turned to a smooth finish. The mould was bored through, so that it could be placed onto the negative pattern for the stamp that had been prepared by the photolithographic process described above. The mould was then filled to its upper edge with PDMS and the threaded holding fixture pressed into the liquid polymer (Fig. 1E). Brushing the inside of the mould with a small amount of 10% SDS facilitated removing the stamps after the PDMS was fully cured. The masters bearing the PDMS filled moulds were kept at room temperature for 48 hours to allow the mixture to polymerize. Faster curing at higher temperatures resulted in distortion of the pattern (Brewer et al., 2011). The cured stamps were removed from the moulds and stored in double distilled water until needed.

**Figure 1:**
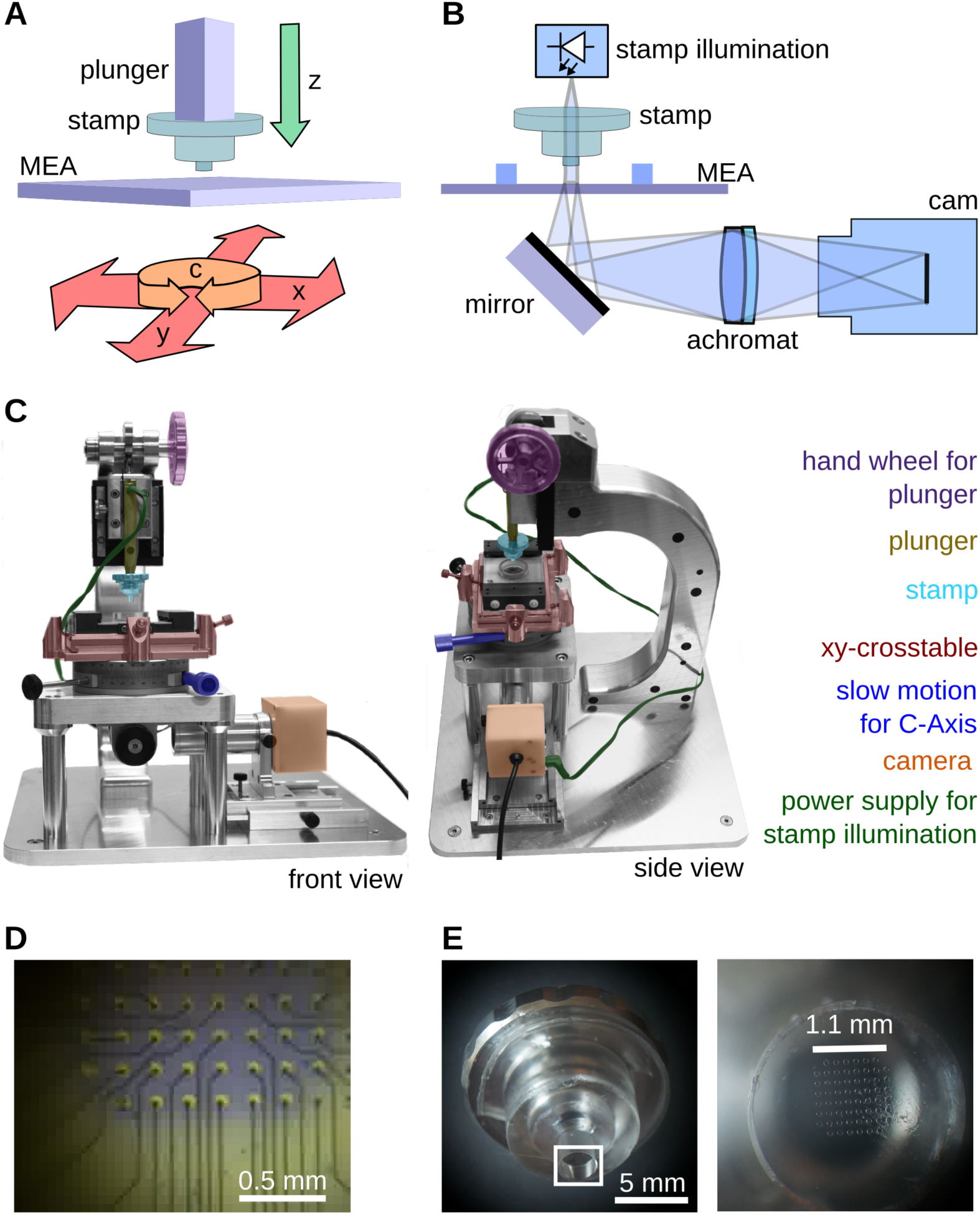
Mechanical and optical setup of the stamping device. The degrees of freedom of the device are distributed as follows: (A) The stamp can be moved in z-direction, whereas the MEA can be translated in the xy-plane, and rotated around the C-axis (B) Optical setup: The transparent stamp is illuminated from above. The positioning of the stamp can be controlled by means of a webcam, onto which the MEA is imaged by means of an achromatic lens L. (C) Photograph of the stamping device. (D) The stamp, as seen from below through a MEA by the webcam. For clarity, a drop of blue ink has been placed on the electrodes. The stamp pattern is in even contact over all sites (E) Micrograph of the stamps and the mould.

**Figure 2:**
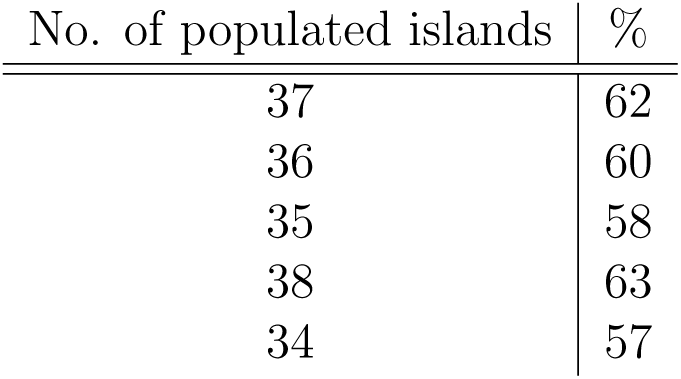
Percentage of populated islands on the 60-Island pattern. Percentage of populated islands of the island pattern consisting of 60 islands aligned to a standard MEA layout (60 electrodes) after 7 days in vitro (n=5; *μ* = 60%; *σ* = ±3.9%).

### 3.4 mechanical setup

The stamp is cast in an aluminium mould that allows for maintaining parallelism of the micro stamp surfaces and the locating face of an aluminium holder moulded into the stamp. This face presses against the contact surface machined onto the plunger. Provided the abutment is kept clean, the surface of the stamps can be kept normal to the axis of the plunger within several seconds of arc. Therefore, it is unnecessary to provide rotational degrees of freedom around the A, and B axes in the stamping mechanism for adjustment.

The stamping press, however, must provide adjustments for the remaining four degrees of freedom: First, the MEA must be rotated around the C-axis, so that the electrode field can be brought into the same orientation as the grid of micro stamps. Then, the MEA must be movable in the x-y-plane to align the electrode field with the stamps (Fig. 1A). The stamp must be depressed by the plunger far enough for the micro stamps to have uniform contact all over the electrode field of the MEA but only so far as not to squash the pattern. This means that the end position of the stamp has to be adjusted to an accuracy of 10 µm. Otherwise, the pattern will be depressed so far that the micro stamps undergo an elastic deformation, similar to Eulerian buckling, which results in instantaneous contact between the MEA and the stamp substrate. To this end, a stopping screw of suitably fine pitch was used to regulate the final position of the stamp relative to the MEA surface. The mechanical setup thus consists of a micrometric cross-table, allowing for positioning of the MEA within a few μm in the x-y-plain. This stage is mounted on a rotary stage with fine movement, providing a rotary degree of freedom along the C axis.

The stamp is fixed to another linear stage with cross-roller bearings, allowing for precise movement in the z-axis. The stamp consists of a clear block of Sylgard 184, cast onto a threaded stamp-holder, the back of which is threaded to allow fixing it to the stamp plunge. The back of this holder also has an accurate locating face matching a similar face on the plunger. The stamp piston is actuated by an eccentric acting against a roller-bearing, fixed to the slide of a vertical stage. The plunger is held in a dovetail fixed to this slide. It is being held there by a small magnet and can be clamped by means of a small screw. This arrangement allows for some limited travel between the slide and the actual plunger, at the same time ensuring that the orientation of the plunger is kept as vertical as possible. All adjustments can be monitored by an inverted microscope built into the apparatus, consisting of an achromat (f=18.5 mm, f/1.5) below the MEA, and a webcam fixed to the side of the mechanism, (see Fig. 1B). Since the stamp material is transparent, the stamp piston and the aluminium have been bored out and an LED has been fixed into the body of the plunger, providing enough light for the contact of the stamp with the MEA surface to be easily recognized, (Fig. 1D). A computer rendering of the stamping machine is included along with the exact design of all parts of the setup and a list of components on a website at http://www.nld.ds.mpg.de/~manuel/website/index.html.

The cost of the setup is small in terms of materials used as many of the elements can be harvested from available old machines or components. Here, we used a cheap commercial webcam for the inverted microscope and the C-Axis manipulator has been taken from a microscopy stage. It is important to note that the specific choice of the manipulators, the webcam or the dichroic is not important as long as the other machine components are modified accordingly. Publishing the construction sheets along with this paper should allow adjusting our setup according to the materials available.

### 3.5 substrate & stamp surface preparation

Sterile MEAs (60MEA200/30iR-Ti; Multi Channel Systems, Reutlingen, Germany) were incubated in purified foetal calf serum (FCS; Gibco) for 30 minutes and washed once with double distilled water and left to dry. A solution of 1% 3-glycidoxypropyltrimethoxysilane (3-GPS (Nam et al., 2004); Sigma Aldrich, Taufkirchen, Germany) in Toluene was added to the culture chamber of the MEA for 20 minutes and subsequently washed three times with Toluene. MEAs were then baked at 100°C for 1 hour and left to cool down to room temperature under a laminar flow hood prior to stamping. Subsequently the prepared PDMS stamps were taken out of the double distilled water and any remaining water was removed by suction. Stamps were sterilized by dipping in 70% EtOH for 10 seconds. Excess EtOH was removed by suction and stamps were left to dry for 5 minutes. A drop of the anionic detergent sodium dodecyl sulphate (SDS; 10% w/v; Sigma Aldrich-Aldrich, Taufkirchen, Germany) was added for 20 minutes on top of the patterned side of the stamp. This adds a release layer between the PDMS and the adhesion promoters to improve transfer to the glass surface, leading to enhanced cellular growth on micro-stamped substrates and increasing the durability of the PDMS stamp (Chang et al., 2002). The SDS was dried under a nitrogen stream, washed with double distilled water and dried with nitrogen again. Aliquots of a 0.1 mg/ml solution of poly-L-lysine conjugated with the fluorescent label fluoresceinisothiocyanat (PLL-FITC) (70,000-150,000 MW; Sigma Aldrich, Taufkirchen, Germany) in phosphate-buffered saline (pH 7.4) were thawed for 1 hour at 37°C in a water bath and vigorously shaken by hand every 15 minutes to dissolve clusters of coagulated PLL-FITC (Brewer et al., 2011). 50 µl droplets of the solution were added to the patterned side of the stamps and incubated in the dark for one hour. Excess liquid was removed and the stamps were left to dry for 10 minutes to allow all moisture to evaporate. It was critical to the stamping process not to allow the PLL-FITC droplet to evaporate before removing it by suction. The stainless steel moulds containing the stamps were then placed into the plunger of the mechanical stamping device, aligned with the MEA and stamped unto the substrate.

### 3.6 cell culture

Cell cultures were prepared according to Brewer (Brewer et al., 1993). Hippocampal neurons were obtained from Wisteria WU rat embryos at 18 days of gestation (E18). The pregnant rat was anaesthetized by CO2. The embryos were removed by a caesarean section, decapitated and transferred to cooled petri dishes. The skull cavity was opened and the brain removed. Hippocampi were surgically extracted and transferred to a HEPES (Invitrogen, Germany) buffer. The supernatant was removed and the extracted hippocampi were trypsinized in a Trypsin/Ethylenediaminetetraacetic acid (EDTA) (trypsin: 0.05%; EDTA: 0.02%; Sigma Aldrich, Taufkirchen, Germany) buffer for 15 minutes at 37°C. Trypsinized cells were then transferred to a 10% Foetal calf serum (FCS) solution. Thorough trituration using a syringe and a needle with a diameter of 1 mm followed. The cell suspension was then centrifuged at 1200 rpm for 2 minutes. The pellet was resuspended in 2 ml of serum-free B27/Neurobasal (B27:5%;Gibco) medium supplemented with 0.5 mM glutamine and Basic Fibroblast Growth Factor (bFGF). Cells were counted with a Neubauer improved counting chamber. A droplet of ≈ 100 µl cell suspension containing 50.000 cells/ml was added on top of the electrode field of the MEAs. This density was chosen to prevent the formation of neuronal cell clusters, as deterioration of the pattern was observed at higher densities. Lower densities led to lower survivability of neurons after more than 7 days in vitro. The MEAs were then kept in an incubator providing a humidified atmosphere containing 5% CO_2_ at 37°C for 4 hours to allow the cells to settle. 1 ml of the B27/Neurobasal medium was then added to the cell chamber. Medium was changed every seven days. All animals were kept and bred in the animal house of the Max-Planck-Institute of Experimental Medicine according to European and German guidelines for experimental animals. Animal experiments were carried out with authorization of the responsible federal state authority.

### 3.7 immunocytochemistry

Patterned cultures on MEAs were used for immunocytochemistry after 14 days in vitro. Cultures were fixed with 4% paraformaldehyde in phosphate buffer (pH 7.4) for 3 minutes at 4°C and subsequently washed three times with phosphate-buffered saline (PBS). Unspecific binding sites were blocked by incubation with 3% Albumin PBS for 30 min at room temperature before cells were permeabilized with Triton X-100 (0.5% in PBS, 5 min, 4°C). Primary antibodies (mouse monoclonal anti-Neurofilament (Abcam, 1:100), rabbit monoclonal anti-GFAP (Abcam, 1:100)) were diluted in 3% bovine serum albumin (BSA) and 0.1% Tween-20 in PBS and then applied overnight at 4°C. After rinsing with PBS, secondary antibodies from donkey were applied for 2 h with a dilution of 1:1000 (alexa 647 anti-mouse IgG, alexa 488 anti-rabbit (Abcam)). MEAs were sealed with round coverslips (15 mm diameter) and mounting medium (ProLong Gold, Invitrogen, Carlsbad, USA), and samples were imaged by fluorescence microscopy using a Zeiss Axiovert 200 (Zeiss, Göttingen, Germany) with a 20x objective.

### 3.8 statistics

Cultures growing on patterned islands were assessed by phase contrast microscopy after 7 days in vitro. Every island populated by at least 1 neuron was counted as populated. Interconnected islands showing more than one interconnection were omitted from the dataset. Mean and standard deviation of the percentage of patterned islands populated by neurons in a sample of X cultures are shown in Table 1.

### 3.9 multielectrode array recording

Recordings were made on a 60 channel MEA amplifier (MEA-1060 Inv, Multichannel Systems, Reutlingen, Germany). Data from MEAs were registered at 25 kHz using a 64-channel A/D converter and MC_Rack software (Multichannel Systems, Reutlingen, Germany). After high pass filtering (Butterworth second order, 100 Hz) events were detected in a cutout recorded 1 ms before and 2 ms after crossing a threshold of -4*σ* of the filtered electrode signal. The threshold was evaluated for every channel individually and typically around -16 μV, The identified events were then aligned at threshold-crossing and averaged. All data analysis was performed in Python.

## RESULTS

### 4.1 island pattern requirements

To demonstrate our technique and the functionality of the mechanical patterning device we chose an island pattern (Fig. 3A). Sixtyfour circles with a diameter of 90 μm were designed to fit on the electrodes of a 60-Channel MEA layout. This pattern allows for easy alignment and quick assessment of cell growth. The diameter and spacing of the islands/electrodes was chosen in order to minimize overlap of the populated islands, while simultaneously allowing growth and adhesion of several neurons per island. The quality of the produced masters was assessed by a Scanning electron microscopy (SEM) (Fig. 3B-C). The negative pattern produced by the photo-lithographic process was checked for accuracy of island diameter and freedom from distortion. The structures seen in the figures represent holes in the photo resist layer. During stamp casting, the PDMS flows into these holes and forms micro-pillars. As printing the coated micro-pillars on the prepared substrate forms the island pattern, it is crucial to assess the quality of photoresist development before using the master as a mould for stamp casting. The steepness of the edges translates into structural integrity of the PDMS micro pillars. This, in turn, affects the amount of pressure the pattern field can tolerate before collapsing during stamping. At the basis of the structures seen in Fig. 3B & C, the exposed surface of the soda-lime substrate has no apparent residues of photo resist. This is important, because incomplete development of the photo resist layer leads to an uneven stamp surface and also uneven printing of the adhesion promoting molecules to the MEA substrate. After stamp casting and substrate preparation according to the protocol described above, the PLL-FITC coated stamps were printed on a MEA substrate and assessed for complete application of the islands pattern on the substrate (Fig. 3D & F). The patterning and alignment made possible by the mechanical patterning device described here lead to faithful, reproducible and highly controllable patterning of the desired topology. Fluorescence imaging showed that all sixty islands were printed on the electrodes without visible distortions or deformations of the pattern. The coating was smooth and relatively homogeneous over all islands. In Fig. 4 illustrates that islands were coated with a variable degree of coating as measured by the fluorescence intensity. The variability between islands seems not to be affected by stamping rounds and is most likely a related to inhomogeneity of the protein coating on the PDMS columns or inhomogeneity of the chemical modification of the glass surface.

**Figure 3:**
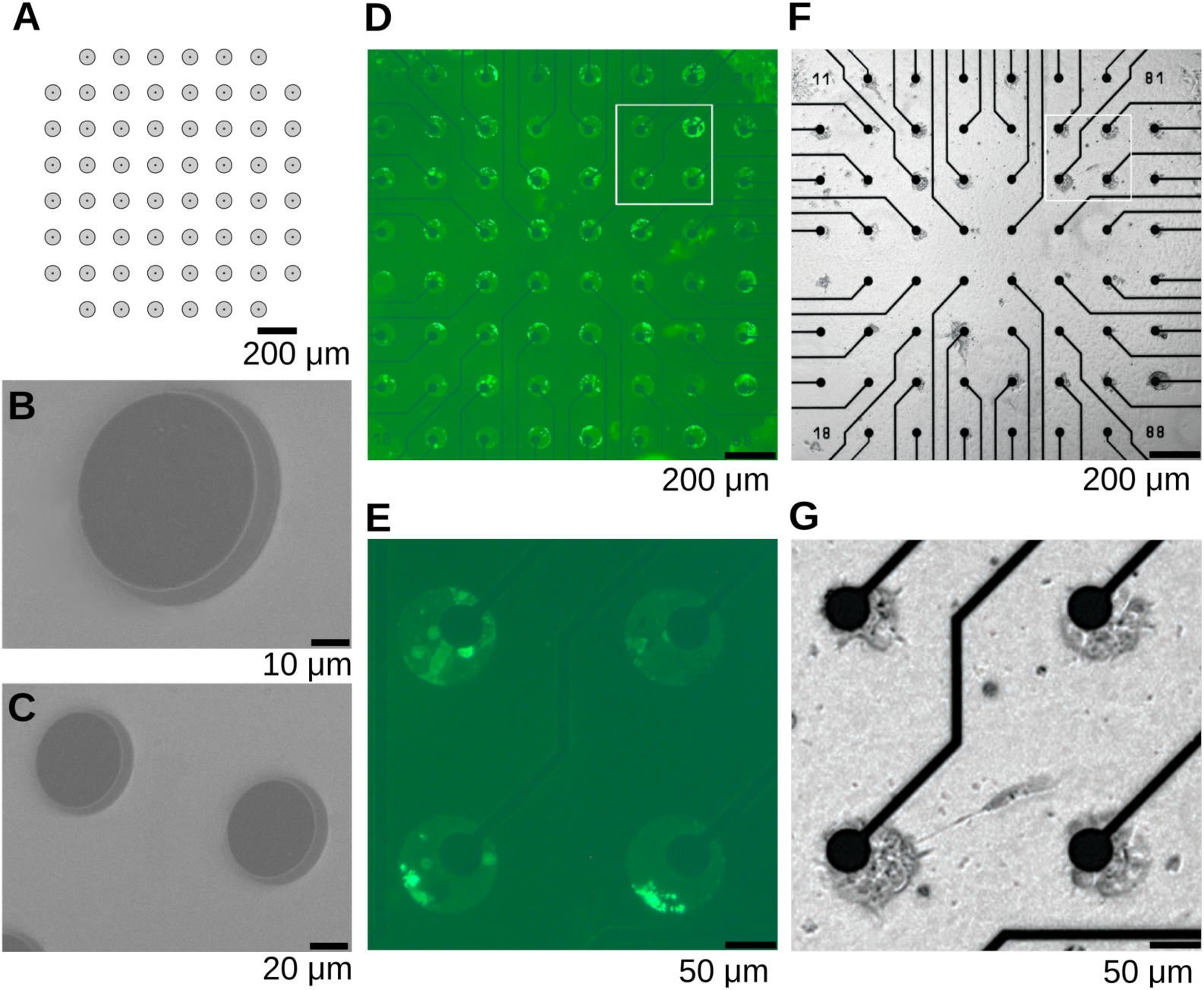
Patterning procedure validation. (A) CAD schematic of the Island pattern: 60 Islands cover the same number of electrodes. Diameter of Islands: 90 μm. Dots represent electrodes. (B – C) Quality of produced masters was assessed by scanning electron microscopy (SEM). Examination focused on the steepness of the edges, indicating a thorough development of the exposed photoresist. The islands in this image are dissolved photo resist holes, showing the soda-lime surface of the substrate at its bottom surrounded by an intact layer of photo resist. (D) Fluorescence of PLL-FITC on aligned and stamped MEA substrate. (E) Higher magnification of (D), the part magnified is marked with a square. Scale bar: 50 pm. (F) Neuronal islands on MEA after 21 DIV. (G) Higher magnification of (F), the magnified part is marked with a square.

**Figure 4:**
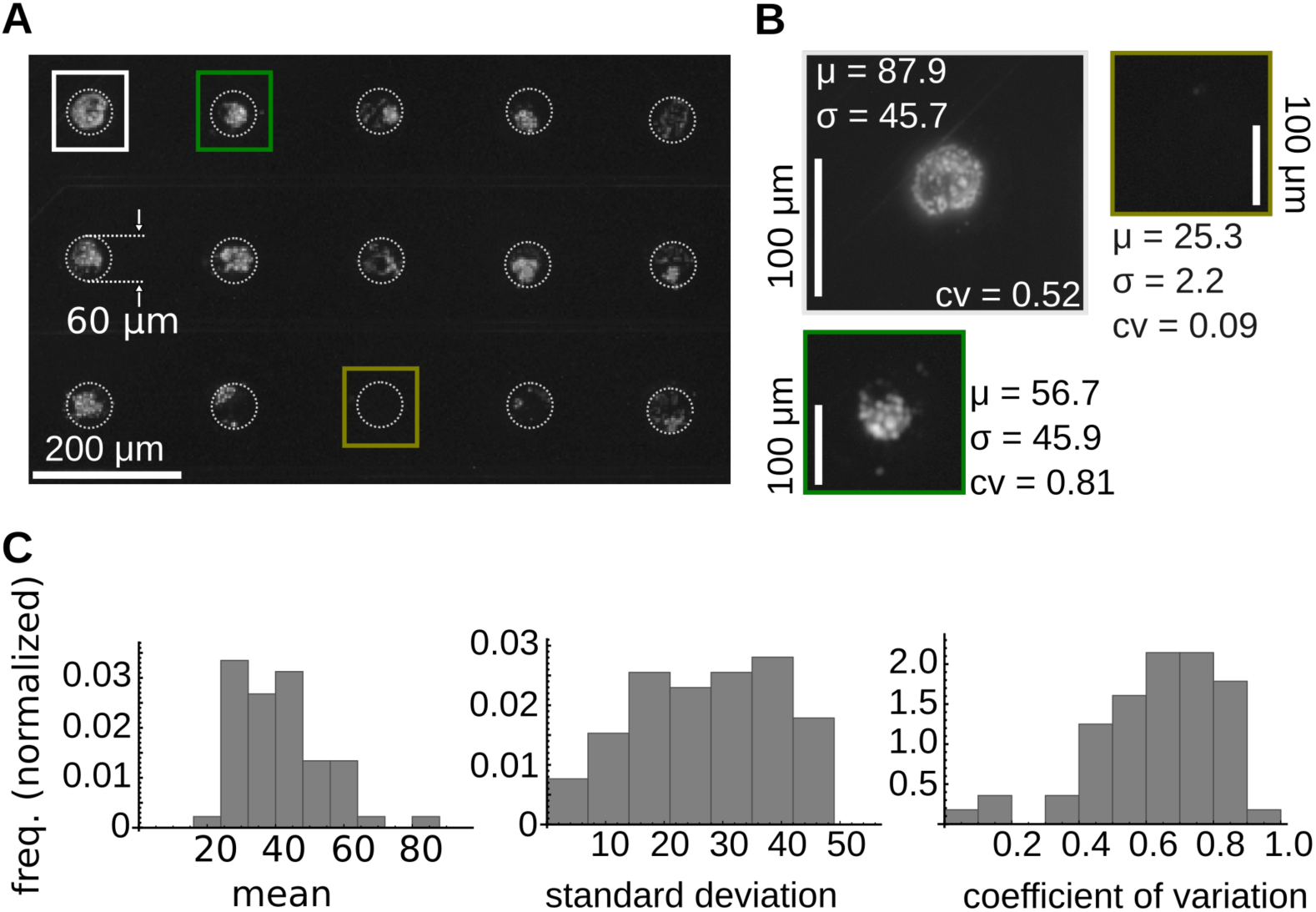
Reproducibility of the patterning procedure. (A) PLL-FITC coated islands stamped on a substrate. The islands have a 60 μm diameter. (B) Magnified PLL-FITC coated islands from (A) along with the mean, standard deviation and coefficient of variation of the fluorescence intensity of the FITC. (C) The histogram of distributions of the mean, standard deviation and coefficient of variation of fluorescence intensity.

**Figure 5:**
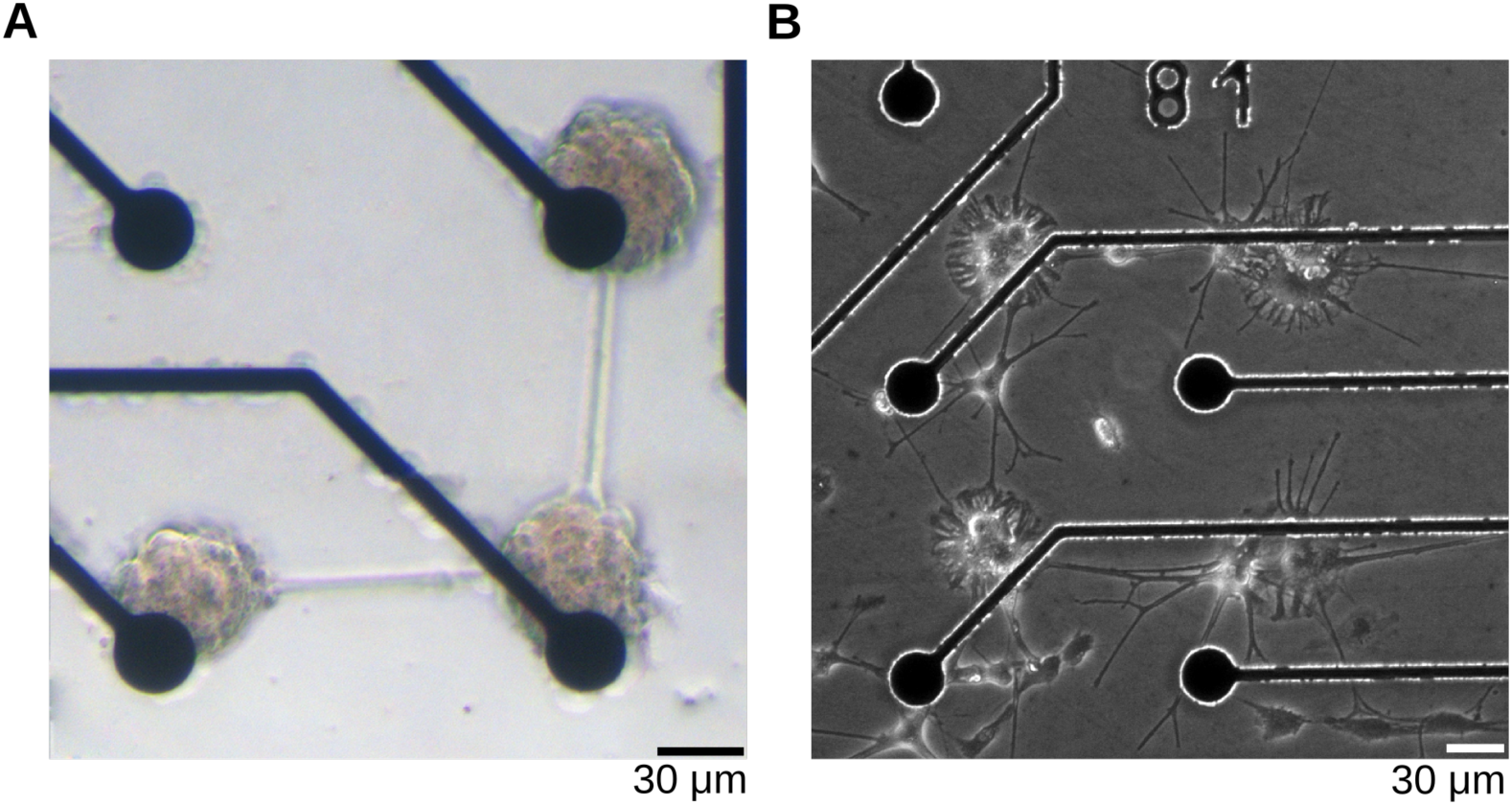
Patterned neuronal cultures. (A) Neuronal islands on the electrodes of the MEA after 14 DIV. (B) Neuronal islands growing beside the MEA electrodes. Patterns were slightly displaced to the electrodes to show that neurons prefer to grow on patterned PLL-FITC over unpatterned electrodes.

**Figure 6:**
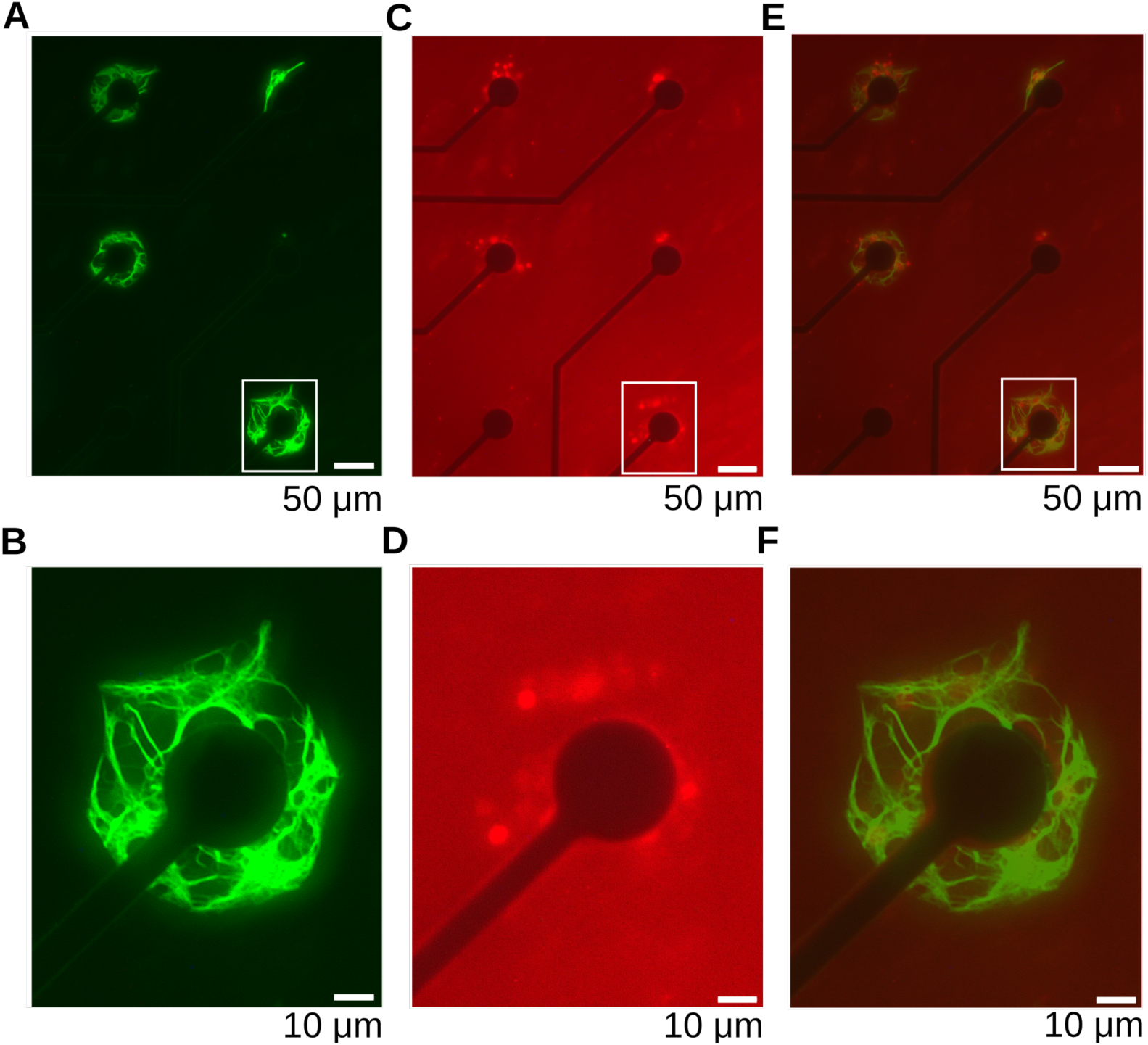
Immunocytochemical staining of patterned neuronal cultures. (A) Neurofilament staining of neuronal islands on MEA. (B) Enlarged from the white frame in (A) Scale bar: 10 pm. (C) GFAP staining of neuronal islands on MEA. (D) Enlarged from the white frame in (C). (E) Overlay showing Neurofilament and GFAP staining of neuronal islands on MEA. (F) Enlarged from the white frame in (E).

### 4.2 neuronal cultures on patterned islands

Hippocampal neurons populated the Islands and showed normal neurite outgrowth (Fig. 3F-G). Cell bodies and processes generally avoided uncoated areas, though some axons reached neighbouring islands (Fig. 4A). Using the AP-μCP technique, the pattern on the stamp can easily be aligned to any feature of the substrate, as shown in Fig. 4B, in which hippocampal neurons are located on islands patterned exactly between the electrodes. Five MEAs were AP-μCP treated with the 64-islands pattern. The percentage of populated islands on the printed and aligned 60-island layout shows that after 7 days in vitro on average 36 out of 60 (60%, n=5; standard deviation= ± 3.9%) islands were populated by one or more neuronal cell bodies and their neurites (Table 1). These results show that the presented patterning method, together with the custom made mechanical stamping device (AP-μCP) is well suited for patterning neurons in a accurate and reproducible manner.

### 4.3 characterization of patterned neuronal cultures by immunocytochemical staining

To determine whether the neuronal cultures populating the islands patterns showed characteristics of typical neuronal networks grown in vitro, neurite structure and astroglial growth were assessed by immunocytochemical staining after 14 days in vitro. Staining against neurofilament, a protein found in intermediate filaments in the axons of neurons, labels axonal processes indicating, that neuronal wiring occurs (Fig. 3A & B). Single neurons growing on the patterned islands enable using the island pattern in studies of autaptic neurons (Bekkers et al., 1991; Pyott et al., 2002). Astrocytes are known to play a substantial role in neuronal development i.e. in metabolic support of neurons, synaptic efficiency and long-term potentiation (Tsacopoulos et al., 1996; Pfrieger et al., 1997; Henneberger et al., 2010). Islands were stained after 14 days in vitro against glial fibrillary acidic protein (GFAP), an intermediate filament of the astroglial cytoskeleton. Each island populated by neurons as marked by staining against neurofilament and thus was accompanied by supporting astrocytes (Fig. 3E & F).

### 4.4 electrical activity in patterned cultures

In order to check the usability of these patterned neuronal cultures for electrophysiological studies, we measured the electrical activity in patterned cultures (Fig. 7A). We found that cultures were showing the stereotypical bursting behaviour observed in in-vitro neuronal cultures (Fig. 7B).The recorded voltage wave forms had relatively large amplitudes on multiple channel along the patterned one dimensional culture. The recorded electrical activity confirms the usability of these patterned cultures for electrophysiological measurements as it demonstrates that the substrate modification did not affect the electrode impedance.

**Figure 7:**
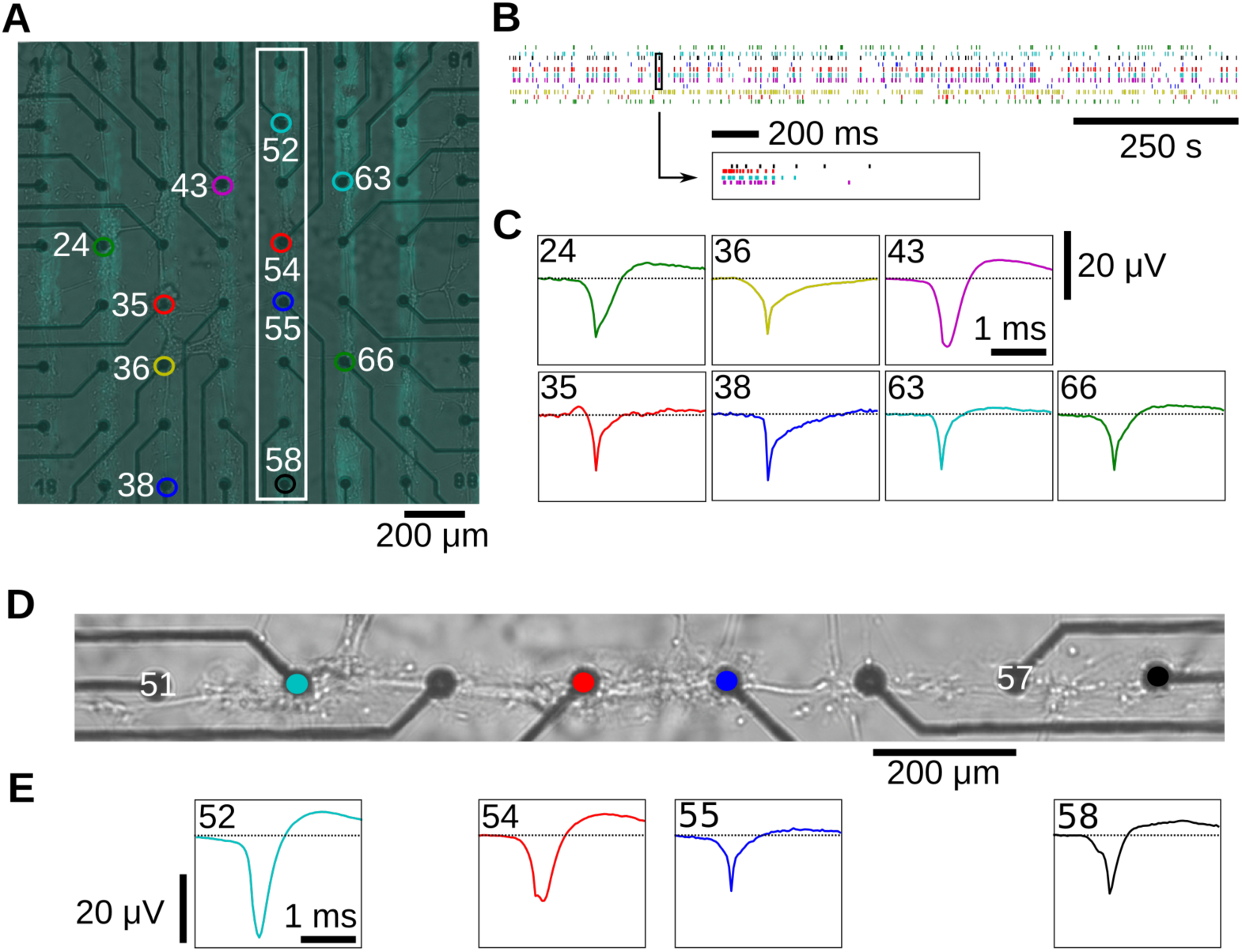
Electrical activity propagation in patterned cultures. (A) The “Highway” pattern of a one dimensional neuronal culture on a multielectrode array. (B) The spike trains of neurons spanning several electrodes. The inset shows bursting activity on a time scale of ms. (C) The waveforms recorded at the electrodes 24, 36, 43, 35, 38, 63, 66. (D) Magnified neuronal line from (A). (E) The waveforms are from electrodes 52, 54, 55, 58.

## DISCUSSION

We presented a novel technique (AP-μCP) that enables reproducible, precise patterning of neurons on MEAs. We have shown that it can produce patterns of high reproducibility while neuronal growth remains intact. AP-μCP achieved a marked and significant increase in pattern coverage, compared to other already established methods. The percentage of populated islands on the printed and aligned 60-island layout in our study shows that after 7 days in vitro, on average 36 out of 60 (60%, n=5; standard deviation= ± 3.9%) islands are populated by one or more neuronal body and its processes. To compare these numbers with the literature, previous studies applying micro-contact printing techniques on MEAs were taken into consideration (James et al., 2000; Chang et al., 2003; James et al., 2004; Heller et al., 2005; Boehler et al., 2012). Few studies (Jungblut et al., 2009; Jun et al., 2007) explicitly state the percentage of pattern coverage over the standard MEA layout: ≈ 47% and ≈ 25% of electrodes covered in average respectively. The average coverage of 60% achieved by AP-μCP thus suggests a significant improvement upon previous methods. The coating is placed on all islands and despite the variability in the coating homogeneity, the neurons seem insensitive to the homogeneity and occupy the majority of islands (60%). Nevertheless, we expect that the method can be further improved by increasing the homogeneity and reproducibility of the coating.

The electrode impedance did not change substantially as evidenced by the recorded electrical activity that showed the hallmarks of in-vitro spontaneous activity highlighting the feasibility of using these patterned cultures for electrophysiological recording. We also reproduced the observation of propagating activity in one dimensional cultures that has been demonstrated previously by Ca imaging (Feinerman et al. 2005).

Patterning of neuronal cultures is regarded as a promising tool to address questions concerning network dynamics and signal propagation. First devised in the 1960’s and 1970’s, topographical patterning methods on planar substrates were achieved by etched or scribed grooves. These first patterning experiments were hard to reproduce, lacking exact means and specifications for placement and alignment. Since then, lithographic procedures have been continuously refined during the past decade (Niemeyer et al., 2004). Micron-sized arrays of silicon pillars were used to guide the growth of neurons and astrocytes (Turner et al., 2000). Electron beam lithography has been used to define features in the nanometer scale for neuronal cell patterning (St John et al., 1997; Curtis et al., 1998), requiring highly specialized equipment. Another way to direct cell growth along topographical substrate modification is by immobilizing cells in well-like structures. Lithographic methods developed for the production of Micro–Electro–Mechanical Systems (MEMS) enable the realization of these features with a good spatial resolution. Neuronal cells were grown at the bottom of deep pits on a silicon substrate and recorded through metal micro-electrodes (Maher et al., 1999). Three-dimensional (3D) microfluidic arrays of poly-dimethylsiloxane (PDMS) were also used to confine cell topology to a certain pattern and were directly structured on silicon wafers using a negative photo resist (Degenaar et al., 2001). Building on the results of these experiments, defined networks of cultured neurons from the pond snail Lymnaea stagnalis have been grown in micro-structured polyester photo resist on a silicon substrate to study interconnected nerve cell pairs using electrophysiological methods (Jenkner et al., 2001). All of these photo-lithographic approaches for physical cell patterning require the continued availability of photo-lithographic facilities and other techniques not generally available in standard cell-culture laboratories.

An alternative approach, chemical patterning methods were introduced in 1965 by adhering fibroblasts to palladium islands evaporated onto a polyacetate surface (Carter et al., 1965). Adhesion promoting molecules can then be transferred to the pattern (Kleinfeld et al., 1988). Due to the techniques involved, organic solvents and alkaline solutions may interfere with the stability of the adhesion promoters. Another method is the patterned deposition of adhesion promoting proteins through silane- or alkanoethiol based surface chemistry (Wheeler et al., 1999; Scholl et al., 2000; Nam et al., 2004). Here, the properties of alkanethiolate monolayers on a substrate are altered through UV-Light exposition, which causes the oxidization to alkanesulfonate, thus altering its solubility. In a second step the exposed area can be linked to a second molecular layer by immersion, creating another monolayer on top of the first (Dulcey et al., 1991). These self-assembled monolayers (SAM) have been used to grow dissociated rat hippocampal neurons on circuit-like patterns (Stenger et al., 1998). SAMs of silanethioles have been used likewise to direct cell growth in vitro (Ma et al., 1998). However, this method requires that the photo-lithographic process is run every time a pattern is created, limiting routine applicability for many biological laboratories. Uniformly coated MEA culture chambers with Poly(L-lysine)-grafted-poly(ethylene glycol) (PLL-g-PEG), a polymer that has cell repelling properties, were locally freed from the polymer by electrical programmable desorption and in a multi process step subsequently coated with Poly-L-Lysine to establish a pattern of cell adhesive molecules. This method was developed for printing patterns of alkanethiolates on a gold substrate (Kumar et al., 1994), and has been used afterwards to topographically confine cell growth of neurons on a glass substrate (James et al., 2000; Lauer et al., 2001; Chang et al., 2006; Jun et al., 2007; Jungblut et al., 2009). Previous work focused on confining cells to uniform patterns of rectangular, striped or triangular shape (Ma et al., 1998; Branch et al., 2000; Liu et al., 2000; Thiebau et al., 2002; Vogt et al., 2003; Vogt et al., 2005; James et al., 2004; Heller et al., 2005; Jungblut et al., 2011).

Previous work describing micro-contact printing techniques on MEAs utilizing alignment of the pattern to a given substrate structure either required expensive microscopy precision placers (Jungblut et al., 2009) or used unspecified custom-made devices (Boehler et al., 2011). The mechanical stamping device presented here relies on few components and a small number of optimized parameters, enabling quick and reproducible alignment of a patterned micro-contact stamp to a substrate layout. With AP-μCP, we present a technique, which can be used in any standard cellculture laboratory, without expensive equipment and continued access to photo-lithography laboratories.

## ACKNOWLEDGEMENTS

The authors would like to thank Markus Krohn and his team at the workshop of the Max Planck Institute of Experimental Medicine, Sabine Stolpe and Sabine Kloppner for help with the cell culture, Tureiti Keith for comments on the data analysis and Elisha Moses for discussions. The authors acknowledge the BMBF for funding (Grant number 01GQ0811, 01GQ01005B, 01GQ0922), ZIM grant (KF2710201 DF0), the DFG (SFB 889 and Cluster of Excellence “Nanoscale Microscopy and Molecular Physiology of the Brain”), and VolkswagenStiftug (ZN2632). A Boehringer Ingelheim Fonds PhD fellowship to Manuel Schottdorf is gratefully acknowledged.

## AUTHOR CONTRIBUTIONS

The project was conceived by Fred Wolf, Walter Stühmer, and Christiane Thielemann and co-supervised by Fred Wolf and Walter Stühmer. Robert Samhaber, Ahmed El Hady and Manuel Schottdorf performed the experiments with the help of Andreas Daus and Kai Bröking. All authors examined and discussed the results. The manuscript was written by Robert Samhaber, Ahmed El Hady, Manuel Schottdorf, Kai Bröking, Walter Stühmer and Fred Wolf.

